# Disease modulation by TIV vaccination during secondary pneumococcal infections in influenza-infected mice

**DOI:** 10.1101/2025.10.16.682853

**Authors:** Juan García-Bernalt Diego, Javier Arranz-Herrero, Gabriel Laghlali, Eleanor Burgess, Seok-Chan Park, Gagandeep Singh, Lauren A. Chang, Prajakta Warang, Moataz Noureddine, Jordi Ochando, Estanislao Nistal-Villan, Michael Schotsaert

**Affiliations:** Department of Microbiology, Icahn School of Medicine at Mount Sinai, New York New York; Global Health and Emerging Pathogens Institute, Icahn School of Medicine at Mount Sinai, New York, NY 10029, USA; Transplant Immunology Unit, National Center of Microbiology, Instituto de Salud Carlos III, Madrid Spain; Department of Microbiology, Icahn School of Medicine at Mount Sinai, New York, NY 10029, USA; Microbiology Section, Dpto. CC, Farmacéuticas y de la Salud, Facultad de Farmacia, Universidad San Pablo-CEU, CEU Universities, 28668, Madrid, Spain; Institute of Applied Molecular Medicine-Nemesio Díez (IMMA-ND), Department of Basic Medical Sciences, Facultad de Medicina, Universidad San Pablo-CEU, CEU Universities, Urbanización Montepríncipe, 28660 Boadilla del Monte, Madrid, Spain; Department of Pharmaceutics, Ghent University, Ghent, Belgium; Graduate School of Biomedical Sciences, Icahn School of Medicine at Mount Sinai, New York, New York, United States of America; Department of Oncological Sciences, Icahn School of Medicine at Mount Sinai, New York New York; National Microbiology Center, National Institutes of Health Carlos III, Madrid, Spain; Icahn Genomics Institute, Icahn School of Medicine at Mount Sinai, New York, New York, United States of America; Marc and Jennifer Lipschultz Precision Immunology Institute, Icahn School of Medicine at Mount Sinai, New York, New York, United States of America

**Author notes:** These two authors contributed equally (Juan García-Bernalt Diego, Javier Arranz-Herrero).

**Keywords:** *Streptococcus pneumoniae*, influenza virus, TIV, bacterial coinfection, bacterial superinfection

## Abstract

Secondary bacterial infections can significantly worsen the clinical course of influenza virus infections and are a leading cause of morbidity and mortality during seasonal influenza epidemics. Despite being a vaccine-preventable disease, influenza-related complications from secondary bacterial infections are an important cause of death, particularly among the elderly population. *Streptococcus pneumoniae* (Spn) is the most common agent responsible for influenza-related secondary bacterial infections. Influenza virus vaccination serves as an effective prophylactic strategy for preventing influenza and reducing the burden of influenza-associated pathology, including secondary bacterial infection. However, whether the protective effects of influenza virus vaccination differ in the context of a secondary Spn infection at the level of the host response remains poorly characterized. Here, we present a preclinical mouse model to examine the impact of influenza vaccination in scenarios involving single infections with influenza A virus H1N1 (NC99) or Spn serotype 1; simultaneous infection with both NC99 and Spn (coinfection), or NC99 infection followed by Spn infection seven days later (superinfection). A single dose of trivalent inactivated Influenza vaccine (TIV) is able to decrease infection lethality in both secondary bacterial infection scenarios. Protection is associated with reduction in both viral and bacterial titers, decreased production of pro-inflammatory cytokines, protection of alveolar macrophages, prevention of exacerbated lung neutrophil recruitment, modulation of neutrophil activation status and induction of lung eosinophil recruitment and activation. These findings underscore the importance of influenza vaccination in modulating disease progression and preventing morbidity and mortality associated with secondary bacterial infections.

## INTRODUCTION

Complications arising from bacterial secondary infections significantly worsen the clinical course of influenza infections, contributing to high morbidity and mortality rates, even in countries with highly developed healthcare systems^1–4^. Despite improvements in our understanding of viral and bacterial pathology, along with better healthcare and extended annual influenza vaccination campaigns, influenza-related complications, especially among elderly patients, continue to claim thousands of lives annually^5^. Moreover, estimates suggest that in the context of the last century’s influenza pandemics, more than 40% of deaths were caused by bacterial secondary infections in 1968-69 and 1957-58, and could be as high as 95% in the 1918 pandemic^6^. Pathology of bacterial secondary infections has been associated with complications largely attributed to acute inflammation of the lower respiratory tissue. While causal agents of bacterial secondary infections in influenza patients vary, *Streptococcus pneumoniae* (Spn) infections are the most prevalent, accounting for over 30% of cases^7^.

Influenza virus infections are characterized by their remarkable capacity to elicit a robust immune responses, leading to acute lung inflammation^8–10^. During primary influenza infections, viral replication mainly targets the respiratory epithelium. This results in the recruitment and infiltration of immune cells, which causes direct damage to the host tissue^10,11^. This damage, coupled with the modulation of host immune responses, creates a conducive environment for secondary bacterial infections^11,12^. Understanding the dynamics of concomitant influenza and pneumococcus infections is critical, as they lead to increased disease severity, higher mortality rates, and additional complications in patient management^13,14^. Numerous studies have extensively explored potential mechanisms through which influenza favors bacterial opportunism^12,15–21^. These investigations have illuminated the intricate interplay between influenza and bacterial pathogens, shedding light on phenomena such as epithelial damage^22^, dysregulation of alveolar macrophages and neutrophils^13,23–26^, coupled with alterations in their phagocytic capabilities^27,28^ and recruitment^29^, or the role of cytokines and chemokines such as IL-10^26^ or CXCL2^30^ driving susceptibility and disease of bacterial secondary infections. Still, research characterizing differential outcomes of bacterial secondary infections that occur at different stages of the influenza infection is lacking. Spn infection kinetics in mice upon lethal intranasal challenge leads to high mortality within 2 to 4 days^31^ while influenza infection takes 5 to 7 to reach peak morbidity and mortality^32^, which is coupled with a significant depletion of alveolar macrophages in lung tissue and bronchoalveolar fluid^33^. Thus, it is to be expected that a secondary bacterial infection occurring within the first 48h of the influenza infection (coinfection) presents a different outcome to one occurring 5 to 7 days later (superinfection).

Currently, vaccination is the cornerstone of prevention plans against influenza epidemics. Licensed influenza vaccines contain either inactivated or live attenuated influenza viruses. Most inactivated vaccines consist of split viruses or subunit influenza antigens. The standard trivalent inactivated influenza vaccine (TIV) is a non-adjuvanted subunit vaccine composed of hemagglutinin molecules from H1N1 and H3N2 influenza A subtypes and one influenza B (Victoria or Yamagata clade)^34,35^. Given the variable efficacy of influenza vaccination depending on the year and antigenic match between vaccines and circulating viruses, it is interesting to investigate the extent with which influenza vaccination approaches and efficacy contribute to the prevention of complications related to bacterial secondary infections, which has been partially addressed in surveillance studies^36,37^, but is still poorly characterized mechanistically. We have previously shown that the administration of TIV protects from mortality after *Staphylococcus aureus* (Sa) bacterial superinfection in a mouse preclinical model, resulting in reduced lung bacterial titers, limited alveolar macrophage loss, and reduced infiltration of inflammatory monocytes in the lung^38^. Other research groups have highlighted that the protection against disease during Sa superinfection by influenza vaccines highly depends on the immune skewing of the host. When the vaccine is administered with Th1-inducing adjuvant formulations (MF59 plus CpG), superior protection is achieved than when Th1/2 (MF59) or Th17 inducers (LTK63) are used^39^. Focusing on *Streptococcus*, subcutaneous immunization with formalin-inactivated influenza A vaccine or intranasal immunization with influenza A vaccine and cholera toxin protected more than 75% of mice from death by lethal influenza A-*Streptococcus pyogenes* superinfection^40^.

To address these research gaps, we present in this manuscript a preclinical C57BL/6 mice model evaluating the multifaceted interactions between influenza virus and Spn, with two different infection regimes: (i) coinfection, defined as a simultaneous infection with a sublethal dose of A/NewCaledonia/20/1999 Influenza virus (NC99, H1N1) and Spn ATCC6301® serotype 1 strain, a highly common and invasive strain in children and in young adults^41^; or (ii) superinfection, defined as an initial infection with NC99 followed, one week later, with a Spn infection. Both coinfection and superinfection were characterized by lung viral and bacterial kinetics, as well as immune profiling of lung myeloid populations and cytokine/chemokine production. Additionally, we investigated the potential role of a suboptimal TIV vaccination dose in the protection from morbidity and mortality in the context of bacterial secondary infections, characterizing immune responses from both humoral and cellular perspectives.

## RESULTS

### A single TIV dose significantly decreased murine morbidity and mortality in Spn coinfection and superinfection

First, we investigated the potential differences in disease outcome between Spn coinfections and superinfections, and evaluated if TIV could have a protective role against Spn secondary infections (Figure 1). We established a C57BL/6 mouse model that was vaccinated with a single dose of TIV [equivalent of 3 μg hemagglutinin (HA), one fifth of the human dose] or mock vaccinated with PBS, a similar model to the one we previously established for Sa^38^. This vaccination regime is suboptimal and conducive to influenza breakthrough infections that cause reduced morbidity [approximately 10% bodyweight loss at 7 days post-infection (DPI)] compared to unvaccinated animals (approximately 20% bodyweight loss), providing a model of co- and superinfection with pre-existing influenza virus-specific immunity. 21 days after vaccination, mice were intranasally challenged with either 50µL of PBS, a sublethal dose of NC99 [632 Plaque Forming Units (PFU), equivalent to 0.4 lethal dose 50% (LD_50_)], a sublethal dose of Spn [10^4^ Colony Forming Units (CFU)] or both (superinfection and coinfection). For coinfection, bacterial and viral intranasal challenges were performed simultaneously on the day of infection (0 DPI), whereas for superinfection, mice were challenged with NC99 at 0 DPI and with Spn at 7 DPI (Fig. 1a).

**Figure 1.**
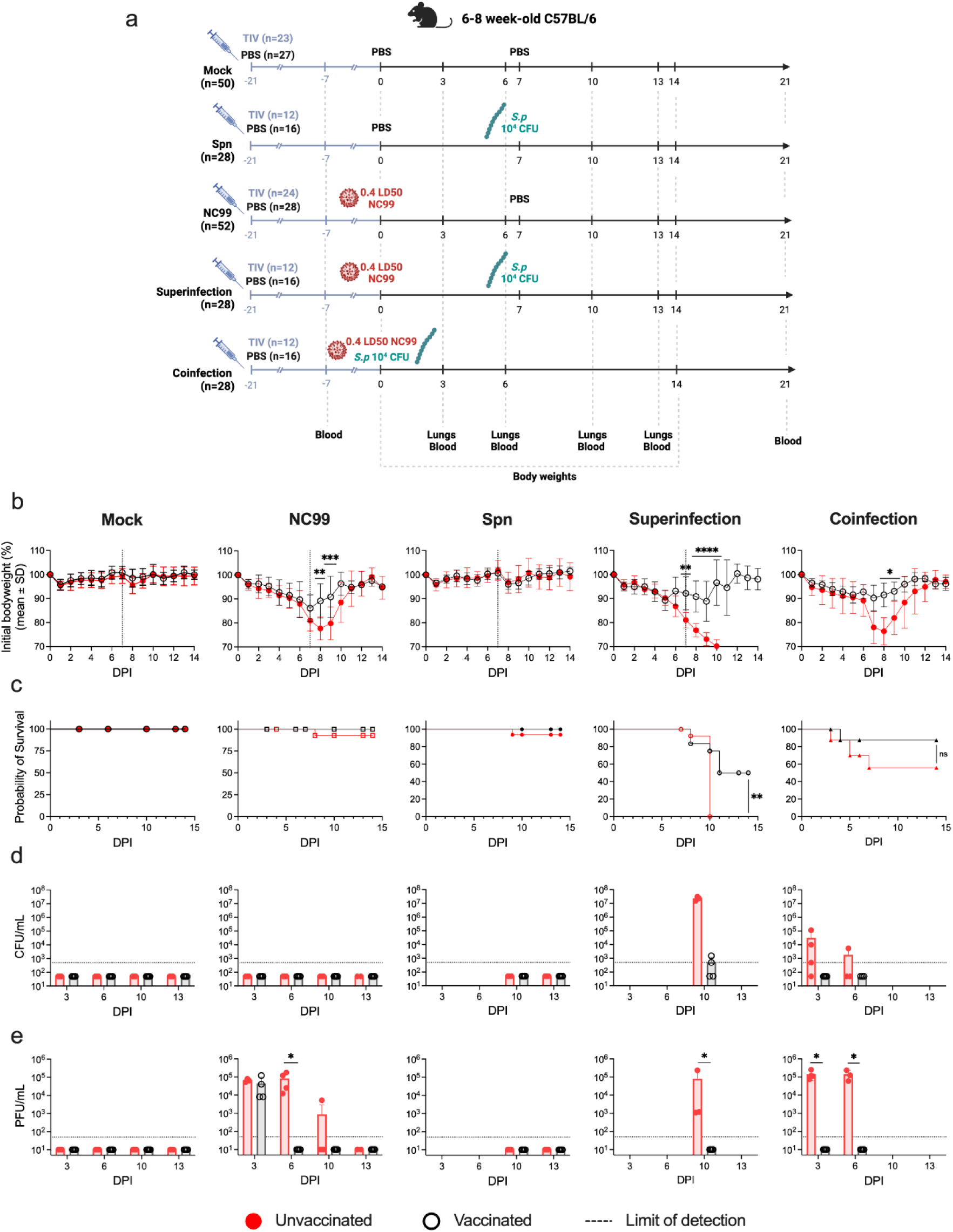
A single TIV vaccination has protective effects against secondary bacterial infections. **(a)** Schematic and graphical representation of experimental timeline. Visual representation of the five conditions: Mock, NC99 infected, Spn infected, superinfection, and coinfection. The timeline highlights days of treatment, infection, and end-point events for sample extraction. Created with: Biorender.com **(b-e)** Panels showing mice bodyweight, survival curves, and lung bacterial and viral titers for the different experimental groups. Female C57BL/6 were vaccinated with TIV (black) or mock vaccinated (red) intranasally inoculated with 50µL of PBS, 0.4LD_50_ of NC99, 10^4^ CFU of Spn, or a combination of infections (coinfection and superinfection). Graphical presentation displaying weight loss **(b)**, mortality, **(c)**, bacterial **(d)** and viral **(e)** titers in the lungs of mice under various conditions. Data from three independent experiments was combined. Differences in bodyweight, bacterial and viral titers assessed by Mann-Whitney U test and differences in survival were assessed by Log Rank (Mantel-Cox) test. Significance values are represented as * p≤0.05, ** p≤0.01, *** p≤0.001, **** p≤0.0001.

As expected, initial infection with NC99 alone induced acute, self-limiting disease, with high morbidity, as reflected by a mean bodyweight loss at 7DPI of 19.02%. In contrast, Spn infection alone caused no significant morbidity (Fig. 1b). While the morbidity associated with coinfection was only slightly higher than NC99 alone (22.03% bodyweight loss at 7DPI), survival rates for coinfected mice decreased considerably compared to NC99 alone (56% survival and 92.5% survival, respectively). Superinfection proved to be more lethal than coinfection with every mouse reaching humane endpoint by 10 DPI, only three days after exposure to Spn (Fig. 1c). Detectable lung bacterial titers were only found in co- and superinfected animals, but not in single Spn infections. Superinfection led to higher lung bacterial titers 3 days after bacterial challenge compared to coinfection (2.29×10^7^ CFUs/mL for superinfection and 3.13×10^4^ CFUs/mL for coinfection) (Fig. 1d). Lung viral titers presented no significant differences between bacterial coinfection or NC99 alone at 3 or 6 DPI. At 10 DPI, while virus titers were cleared in 5 of 6 animals infected only with NC99, the superinfected group still presented high titers (7.91×10^4^ PFUs/mL), underscoring the impairment of efficient antiviral immune responses in the context of a Spn superinfection (Figure 1e).

After vaccination, the mice morbidity and mortality were significantly reduced. Vaccinated mice challenged with NC99 alone lost significantly less bodyweight and started recovering sooner than unvaccinated mice. Similar observations were made in coinfected animals, for which vaccination also improved survival (from 56% to 87.5%). Vaccination also reduced superinfection morbidity and mortality, significantly reducing body weight loss after Spn challenge and improving survival from 0% to 50% (Figures 1b, c). Moreover, lung bacterial growth was controlled as soon as 3 DPI in coinfected mice, and only half of the superinfected mice presented detectable titers at 10DPI (Figure 1d). In line with previous findings for our challenge model, some NC99 breakthrough infection is observed, with animals challenged with NC99 alone presenting comparable lung viral titers at 3DPI in vaccinated and unvaccinated animals (4.4×10^4^ PFUs/mL and 6.5×10^4^ PFUs/mL, respectively) but controlling the infection much faster, with no detectable titers at 6DPI. Strikingly, vaccinated animals that were coinfected cleared viral infection more efficiently than those challenged with NC99 alone, showing no detectable titers already at 3 DPI. Lastly, vaccination resulted in no detectable lung viral or bacterial titers at 10 DPI in superinfected mice (Figure 1e).

### Increased expression of Siglec-F in lung neutrophils and eosinophils correlates with protection from bacterial secondary infections after TIV vaccination

To further characterize the immune response after infection with and without vaccination in the different conditions, lungs were collected on 6 and 10 DPI. Lungs were dissociated into a single cell suspension and relative quantification of immune cell subsets was performed by flow cytometry (Fig. 2 and Fig. S1). Gating strategy is detailed in Fig. S2. Alveolar macrophage dysregulation has been characterized as one of the key mechanisms triggered by influenza infection that promotes increased susceptibility to bacterial secondary infections^33^. Alveolar macrophages exhibited a significant reduction in NC99-challenged mice at 6DPI, which TIV vaccination largely prevented. Interestingly, coinfected animals showed a similar reduction in alveolar macrophages at this stage, however, there was no significant protection of these cells provided by vaccination (Fig. 2a). Infiltration of inflammatory monocytes was significantly increased in both coinfection and NC99 alone groups, but it was prevented by vaccination in both groups (Fig. 2b). At 10 DPI, alveolar macrophages remained low in unvaccinated mice challenged with NC99 alone, similar to superinfected animals, while vaccination showed a trend of alveolar macrophage protection in both groups (Fig. 2a). Interestingly, inflammatory monocytes presented elevated levels in all superinfected animals, regardless of their vaccination status, and significantly higher than the ones in animals infected with NC99 alone or Spn alone (Fig. 2b).

**Figure 2.**
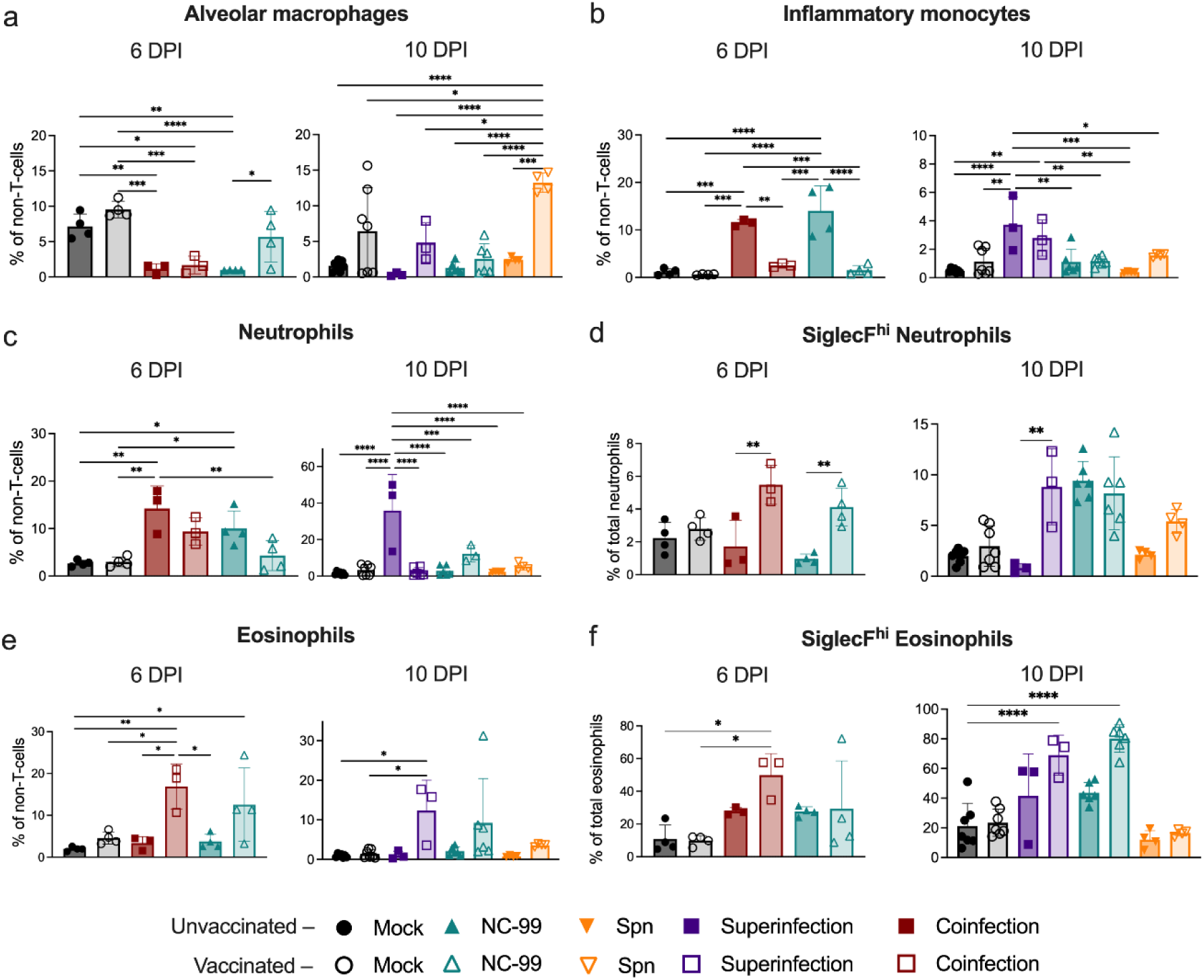
TIV vaccination prevents from exacerbated lung neutrophil recruitment and promotes eosinophil recruitment. Relative quantification of cells identified in lung by flow cytometry. Frequency of (a) alveolar macrophages (b) inflammatory monocytes, (c) neutrophils, (d) SiglecF^hi^ neutrophils, (e) eosinophils and (f) SiglecF^hi^ eosinophils in lungs at 6 and 10 DPI. Comparisons were performed by One-Way ANOVA with Tukey’s multiple comparisons test. Significance values are represented as * p≤0.05, ** p≤0.01, *** p≤0.001, **** p≤0.0001.

Exacerbated lung neutrophilic inflammation in influenza virus-Spn coinfections has been associated with pneumococcal persistence in the lung and increased disease severity^42^. At 6DPI, we observed a significant increase in neutrophil infiltration in NC99-challenged unvaccinated mice (10.02±3.69%), and coinfected mice (14.21±4.75%) compared to the mock unvaccinated condition (2.66±0.65%). A non-significant trend towards the reduction of neutrophilic infiltration by vaccination was observed in mice challenged with NC99 (4.30±3.16%) and coinfected (9.38±2.86%). At 10 DPI, a very strong neutrophil recruitment was observed in unvaccinated superinfected mice, reaching mean values of 35.93±19.64% of total non-T-cells, compared to 1.35±0.97% from unvaccinated mock. Vaccination achieved significant control of neutrophilic infiltration, although neutrophil levels were still elevated (12.22±4.50%) compared to mock or NC99 alone (Fig. 2c). Lung neutrophil subpopulations are still poorly characterized, but they can be assessed based on their expression of Siglec-F. While we observed that neutrophilic lung infiltration was clearly reduced by vaccination, as outlined above (Fig. 2c), the percentage of Siglec-F^hi^ activated neutrophils increased after vaccination in all infected groups. At 6 DPI, both coinfected and NC99 challenged groups that received the vaccine showed a significant increase when compared to the unvaccinated counterparts. Interestingly, at 10 DPI, while TIV vaccination still increased the percentage of Siglec-F^hi^ neutrophils in all infected groups, unvaccinated mice receiving NC99 alone also show an elevated percentage, similar to the mice that received the vaccine (Fig. 2d).

The role of eosinophils in viral infections, especially in the context of vaccination remains a highly debated topic^43^, with both examples of pathological effects mostly associated with vaccine-associated enhanced respiratory disease (VAERD)^44^ and protective effects in the context of breakthrough respiratory infections^45,46^. In our model, an increase in lung eosinophil recruitment was observed for vaccinated groups challenged with NC99 alone (both at 6 and 10 DPI), coinfected (6 DPI) and superinfected (10 DPI), but was absent in all the groups that did not receive the vaccine (Fig. 2e). Similar to neutrophils, lung eosinophils can also be classified based on Siglec-F expression, with Siglec-F^hi^ eosinophils corresponding to the inflammatory eosinophil subset and Siglec-F^int^ eosinophils corresponding to tissue-resident eosinophils. Functionally, Siglec-F^hi^ eosinophils have been associated with exacerbated Th2 responses in murine models for asthma^47,48^. However, previous results from our group and others suggest a protective role of these Siglec-F^hi^ eosinophils in the context of influenza^45,49^. Opposite to the total neutrophil population, total eosinophils increased in vaccinated groups challenged with NC99 alone, coinfected, or superinfected, compared to unvaccinated. In line with our previous findings, the percentage of eosinophils classified as Siglec-F^hi^ significantly augmented in vaccinated animals that were coinfected (6 DPI) and challenged with NC99 alone or superinfected (10 DPI). Unvaccinated mice challenged with NC99 also show an increase in inflammatory eosinophils compared to the mock, although milder than the one in vaccinated groups (Fig. 2f).

Lung CD3^+^ cells did not reveal clear differences among experimental groups. Dendritic cells in the lung increased in vaccinated animals when compared to unvaccinated groups at both 6 DPI and 10 DPI (Fig. S1).

### Cytokine and chemokine profiling of the lungs of superinfected infected mice reveals a differential inflammatory profile

To study cytokines and chemokines associated with the phenotype of the different infection regimes in the lungs of the infected mice, we performed a multiplex-ELISA (Luminex) quantification of lung homogenates supernatants at 3, 6 and 10 DPI (Fig. 3, S3-S5). At 3DPI, mock and Spn infected mice, regardless of their vaccination status, did not show increased levels of any of the cytokines and chemokines measured. Mice infected with NC99 alone showed non-significant increases in the pro-inflammatory chemokines CCL5, CCL7 and CXCL10 compared to mock, that were significantly elevated the vaccinated group. Coinfected animals that did not receive TIV presented a further exacerbated proinflammatory profile, with significantly higher levels of CCL5, CCL7, CXCL10 and IFN-γ. IL-2 was also elevated in all groups challenged with NC99 or coinfected, regardless of their vaccination status. Eosinophil-recruiting eotaxin was significantly elevated in vaccinated groups challenged with NC99 alone or coinfected, in the absence of any of the classical type-2 skewing cytokines (IL-4, IL-5, or IL-13).

**Figure 3.**
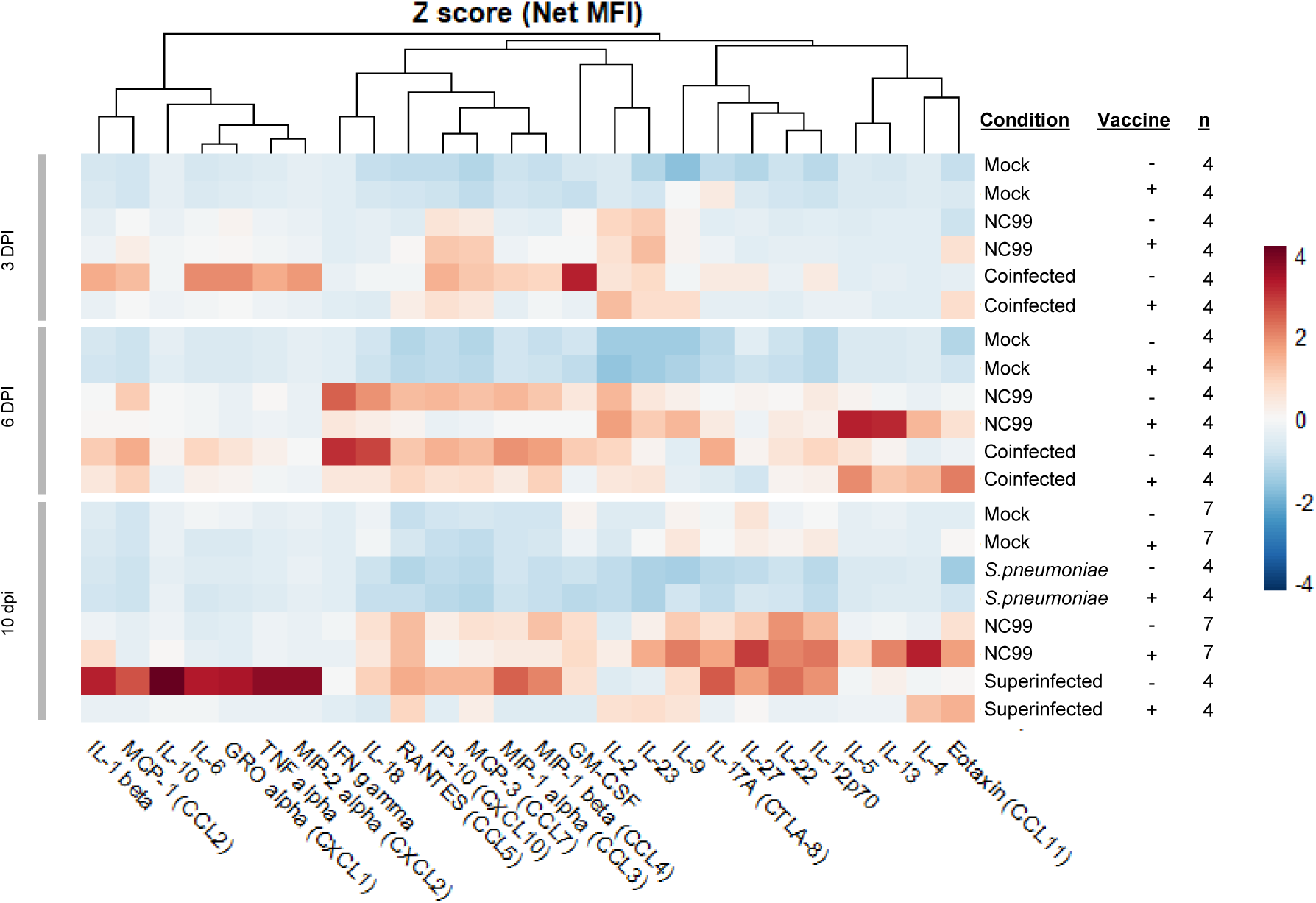
TIV vaccination reduces pro-inflammatory cytokine and chemokine production in lungs from coinfected and superinfected animals. Cytokine profiling from mice lungs at 3, 6, or 10 DPI by Luminex multiplex cytokine ELISA. Heatmap of cytokine and chemokine Z-scored Average Net MFI.

At 6 DPI, differences between vaccinated and unvaccinated mice challenged with NC99 became more apparent, in accordance with the self-limiting nature of the breakthrough infections in the model described above. Chemokines CCL5, CCL7, and CXCL10 remained significantly elevated in unvaccinated mice, while they returned to mock levels or were reduced in vaccinated mice. Unvaccinated coinfected mice presented a more inflammatory cytokine profile at 6DPI than at 3 DPI, with significantly elevated levels of cytokines IL-6, IFN-γ, and IL-18, and chemokines CXCL1, CXCL2, CXCL10, CCL3, CCL4, and CCL7. All these cytokines and chemokines were at the base level or reduced in TIV-vaccinated mice. IL-2 remained elevated in all groups challenged with NC99 or coinfected, regardless of their vaccination status. Eotaxin was not significantly elevated anymore by 6DPI in vaccinated animals challenged with NC99 or coinfected, although non-significant increases in anti-inflammatory cytokines IL-4 and IL-13 could be observed in lungs of vaccinated NC99-infected and coinfected mice at 6 DPI.

Superinfection generated a divergent chemokine and cytokine profile in the lung at 10 DPI, when compared to animals challenged only with NC99. While chemokines such as CCL2, CCL7, and CXCL10 and cytokines such as IL-2 or IFN-γ were still elevated, they trended downwards in NC99 challenged mice, but they were further increased in superinfected animals. Additionally, other cytokines such as IL-6 and IL-10, and chemokines CXCL1 and CXCL2 were highly elevated in the superinfected group. TIV vaccination completely abrogated any increase for most of those cytokines and chemokines, further highlighting the protective role of TIV not only against bacterial proliferation, but also against inflammation.

### A single TIV vaccine dose efficiently induces neutralizing antibodies against NC99, which are boosted by infection, regardless of the secondary bacterial infection status

To evaluate the role of Spn coinfection and superinfection in the humoral response after influenza infection and vaccination, half of the mice included in this study were vaccinated with a single dose of TIV vaccine, and the other half of the mice received a control PBS injection. Two weeks later, TIV-binding IgG titers were measured, and all vaccinated animals showed positive seroconversion (Fig. 4a). The kinetics of the antibody response were also evaluated after the challenge. Given the setup of this experiment with many mice at the beginning, mock-challenged unvaccinated animals showed no detectable TIV-binding IgG, while mock-vaccinated animals showed detectable TIV-binding IgG titers that remained constant from 3 DPI to 21 DPI. Comparable results were obtained in the group that was only challenged with Spn. Challenge with NC99 alone led to the production of detectable TIV-binding IgG titers as soon as 6 DPI in unvaccinated animals. By 21 DPI, vaccinated and unvaccinated animals challenged with NC99 showed comparable IgG titers. While Spn superinfection did not have a significant effect on TIV-binding IgG titers in vaccinated animals, coinfection led to a significant increase in TIV-binding IgG titers when compared to mock-vaccinated animals (p<0.0001), but more interestingly also compared to NC99 challenged vaccinated mice (p<0.0001) (Fig. 4b), which correlates with the faster viral clearance observed in vaccinated coinfected mice. No detectable specific antibodies could be collected for the 21 DPI timepoint for unvaccinated animals in the superinfection group as all mice reached humane endpoint prior to that timepoint.

**Figure 4.**
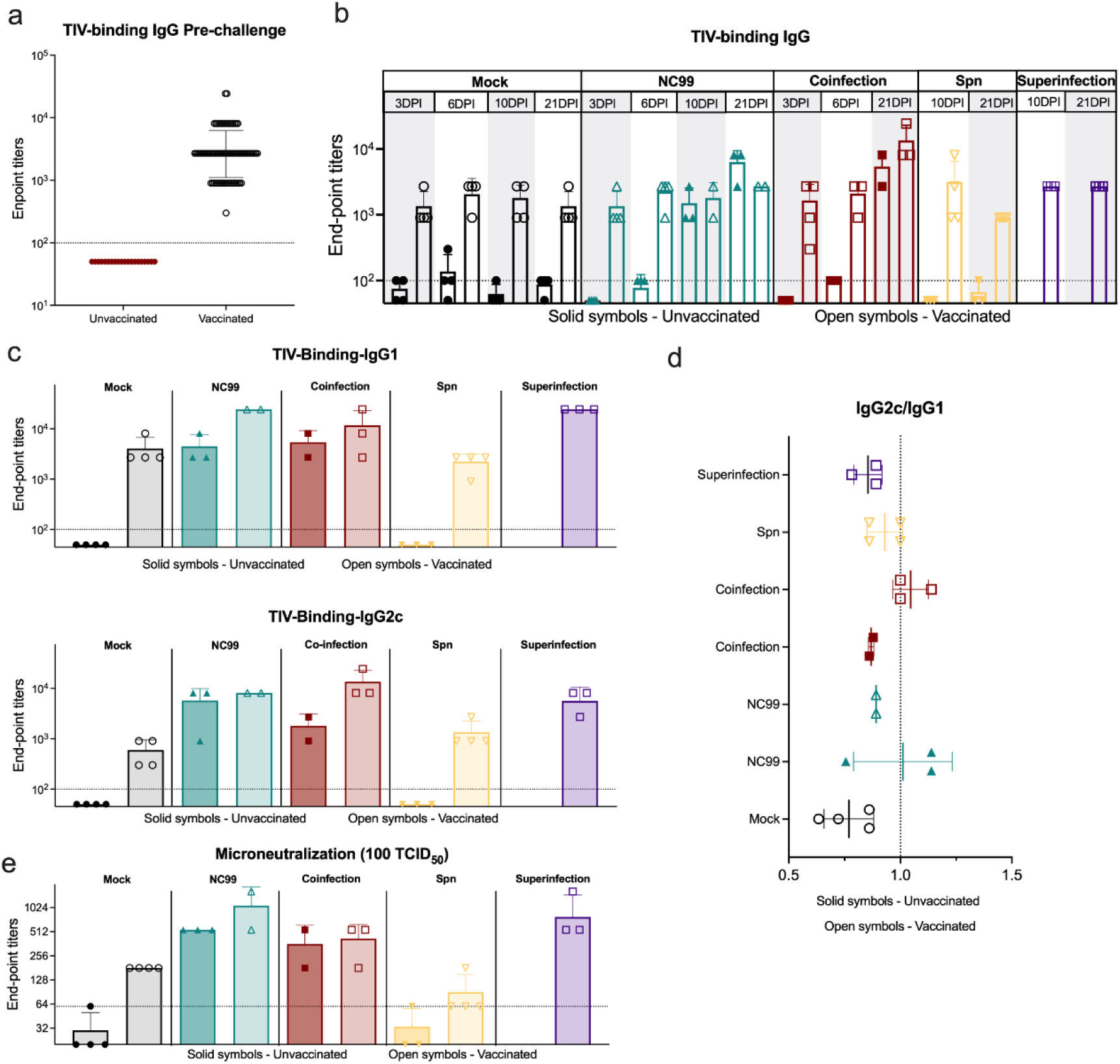
TIV vaccination induces neutralizing antibodies against NC99 with superinfection and coinfection skewing antibody responses. a) TIV-binding IgG endpoint titers two weeks after a single-dose TIV vaccination. b) Kinetics of TIV-binding IgG after challenge, at 3, 6, 10, and 21 DPI. C) TIV-binding IgG1 and IgG2c 21 DPI. D) Coefficient of the log10 of the endpoint titers of TIV-binding IgG2c over the log10 of the endpoint titers of TIV-binding IgG1 (indicative of Th1/Th2 antibody skewing). E) Endpoint titers of the microneutralization assay against 100 TCID_50_ of NC99 in MDCK cells.

Then we evaluated the effect of vaccination and secondary bacterial infection on IgG subtypes. Both IgG1 and IgG2c were boosted by infection 21 DPI in all groups (Fig. 4c). No significant differences were found between vaccinated challenge only with NC99 or coinfected with Spn. Vaccinated superinfected mice showed significantly higher levels of TIV-binding IgG1 compared to vaccinated coinfected mice (p=0.0254) (Fig. 4c). Consequently, vaccinated superinfected mice showed a more Th2-skewed antibody response when compared with mice that were challenged only with NC99 or coinfected (Fig. 4d).

Finally, we performed microneutralization assays to evaluate neutralizing antibody responses against NC99. Vaccinated mice that were superinfected or infected with NC99 alone showed higher microneutralization endpoint titers than vaccinated animals that were coinfected, however, these differences were not significant (Fig. 4e).

In all, our results show that TIV vaccination has a major role in the protection against disease and mortality in influenza infected mice due to secondary bacterial infections, reflected as well in the rate of viral and bacterial clearance, the cellular infiltration in the lung, the cytokine and chemokine profile and the antibody immune response.

## DISCUSSION

Despite the numerous advances in this field, complications derived from Spn secondary bacterial infections in the context of influenza remain a global healthcare problem. Several studies have shed light on the mechanisms underlying the interplay between these two pathogens^17,50–53^. However, many questions remain unanswered. Comparative analyses evaluating secondary bacterial infections based on the timing of the bacterial challenge with respect to the beginning of the influenza infection are scarce. Still, relative timing of the two infections could play a major role in the outcome of the disease due to the differential kinetics of influenza and Spn infections within the host. Here, we distinguished between coinfection (simultaneous infection with both pathogens) and superinfection (infection with Spn 7 days after NC99 challenge), comparing, from an integrative point of view, protection from disease, bacterial and viral clearance and cellular and humoral immune response to influenza virus in the two different scenarios. We also explored the role of pre-existing immunity to influenza, provided by TIV vaccination, on modulation of the host response to infection.

Our results show that mortality rates are elevated in both coinfections and superinfections in the context of NC99 and Spn challenges that are sublethal on their own. Relevant differences were detected between the two secondary infection regimes. Superinfection led to 100% mortality within 3 days of the bacterial challenge, while coinfection showed less than 50% mortality. Therefore, timing of both infections determines the degree of synergism between bacteria and virus to cause a more lethal disease. Other researchers have reported similar observations, where infection with Spn 7 days before influenza infection did not affect the mortality, coinfection at 0 DPI decreased survival rates by almost 50% and if the bacterial infection occurred 3, 5 or 7 days after influenza challenge (superinfection), the lethality reached 100%. Infection with Spn at later timepoints was progressively less lethal^54^. Notwithstanding, these results might be dependent on a plethora of factors, including genetic background of the pre-clinical model or bacterial and viral strains. For example, another study observed extreme susceptibility to Spn in superinfected C57BL/6, the model used in our study, but not in BALB/c^16^.

Additionally, we investigated the impact of vaccination with TIV in our co- and superinfection model. Previously our group has showed the protective capacity of this vaccine against superinfections with *Staphylococcus aureus*^38^. Here we show that even a single dose of TIV, which is considered a suboptimal vaccination regimen as it allows breakthrough influenza infections, significantly increased survival in both superinfections and coinfections, but with different efficacy. This effect was primarily explained by the viral and bacterial titers found in mouse lungs, with lower lung virus titers in vaccinated mice, compared to unvaccinated mice. In unvaccinated NC99 mice, viral clearance was almost complete by 10 DPI. However, unvaccinated superinfected mice showed prolonged virus replication with detectable virus even at 10 DPI. This delay in viral clearance has been previously described and correlated with the initial bacterial inoculum size, among other factors^13^. These results align with what we observed in our experiments and emphasizes the beneficial effects of vaccination in secondary bacterial infections. This is further highlighted by the results obtained in coinfected animals. A single TIV dose is not sufficient to provide sterilizing immunity in mice, as we have previously established in our pre-clinical models^38,45,49^. Here, while breakthrough infection was detectable 3 DPI in vaccinated mice that were challenged only with NC99, coinfected mice were able to clear the virus more efficiently and showed no breakthrough infection at 3 DPI. While the literature describing the effects of influenza infections in the clearance bacterial secondary infections is abundant, the effect of prior or concurrent bacterial infections in viral clearance is less characterized. Early viral control due to prior bacterial infections has been described in other bacteria-virus combinations, such as SARS-CoV-2 and *Mycobacterium tuberculosis*. This control took place as early as one day after viral challenge and was dependent on bacterial dose. The authors speculate that *Mycobacterium*-driven innate inflammation (through PPRs signaling) together with the induction of antiviral interferons, may mediate this early protection, however they are not able to isolate a specific signaling pathway or combination of them responsible for this response^55^. The impact of co-administration of bacteria and virus on the host innate immune response, eventually resulting in better virus control, requires further study.

According to literature, mice that survive Spn infection usually reach bacterial clearance within the third day after infection. On the other hand, if bacterial disease manages to progress, mice typically die between the second and the fourth day after bacterial infection^54^. In line with these observations, in our model, unvaccinated mice infected with only Spn did not present detectable bacteria in lungs at 3 DPI, while in coinfected animals, Spn was found even at 6 DPI.

It is well established that influenza infection decreases macrophages’ phagocytic ability^13^, as well as the absolute number of alveolar macrophages as soon as 3 DPI^50,56^. During the initial week post-influenza infection, over 90% of resident alveolar macrophages are depleted, with the surviving cells exhibiting a necrotic phenotype^50^. After the depletion of alveolar macrophages upon influenza infection, they are repopulated with bone-marrow monocyte-derived alveolar macrophages within days 7-11^56^. These results are in accordance with the extensive macrophage depletion we observed that in both coinfection and superinfection models, that is partially avoided by TIV vaccination. Therefore, a possible explanation for the enhanced mortality if Spn infection occurs after influenza virus infection, is due to the loss of alveolar macrophages that occurs in the days post influenza virus infection^49^, since alveolar macrophages are a first line of defense against bacterial infections, virus-mediated macrophage death may leave mice more susceptible to severe disease outcome during superinfection.

The secondary stimulus for superinfection triggered a massive recruitment of neutrophils. Exacerbated lung neutrophilic infiltration has been widely characterized in superinfection^57^, and it is commonly related to functional impairment in the lungs and lethality. Both coinfected and superinfected mice showed an increase in neutrophil recruitment, that is prevented with a single vaccination dose of TIV. This is paired with the cytokine profile of both conditions, exhibiting elevated values of IL-6, CXCL1, and CXCL2, which induce the production and recruitment of neutrophils. Some studies have focused on neutrophils in superinfections as a major cause of lethality and found protective effects of CXCR1/2 antagonist during Spn superinfections^58^. Once again, TIV vaccination largely prevented excessive neutrophil recruitment to the lung. We further characterized neutrophil populations in the lung, based on their expression of Siglec-F. Dogmatically, Siglec-F expression was typically attributed to eosinophils and alveolar macrophages in the mouse lung. However, a Siglec-F^hi^ lung neutrophil subset has recently been described in the allergy field, characterized by a higher activation and increased effector functions such as neutrophil extracellular trap (NET) release and airway hyperresponsiveness^59,60^. Siglec-F^hi^ neutrophils have also been associated with protection against nasal challenge with *Bordetella pertussis*. Mobilization of this neutrophil subset was mediated by IL-17 and was lost, together with the improved protection, in IL17A^−/−^ mice^61^. In our model, while exacerbated neutrophil recruitment is greatly prevented in TIV vaccinated groups, an increased fraction of Siglec-F^hi^ neutrophils is detected when compared to unvaccinated controls. Therefore, this neutrophil subset could have a role in the observed improved protection against bacterial secondary infections in those groups. Nevertheless, IL-17A concentration in lung supernatants shows very limited differences among groups. Thus, further mechanistic studies are needed to define how these neutrophils are recruited in the context of vaccination and what is their role in protection.

A pronounced lung eosinophilic recruitment was also detected in vaccinated mice challenged with NC99 alone or co/superinfected. Pulmonary eosinophilia is typically associated with negative outcomes of infection in vaccinated individuals, in a phenomenon known as vaccine associated enhancement of respiratory disease (VAERD), initially described in the context of respiratory syncytial virus vaccination^62,63^. However, growing scientific evidence also suggests an antiviral role for eosinophils both mice and humans^64,65^. Our group has previously described similar findings, with vaccine-associated pulmonary infiltration of eosinophils in the influenza mouse and hamster models that rather correlates with protection in the absence of VAERD, in the context of SARS-CoV-2, influenza infection and superinfection with Sa. This pulmonary eosinophilia is vaccine-dose dependent and is present regardless of the immune skewing of the host caused by the vaccine adjuvant^38,43,45,46^.

While we show that a single dose of TIV vaccine is enough to provide detectable TIV-binding IgG titers, virus-specific antibody titers are boosted differently after infection. Notably, vaccinated animals that were coinfected with Spn show significantly higher titers at 21 DPI than animals that received TIV and that were challenged with NC99 alone. Comparatively, superinfected mice, even in the context of vaccination, showed reduced antibody titers compared to coinfected animals. A reduced B-cell response against influenza virus during superinfection is well characterized in the literature, with superinfected mice showing reduced levels of virus-specific IgG, IgM, and IgA, as well as the number of B cells, CD4^+^ T cells, and plasma cells^66^. Superinfections have also been associated with spleen atrophy resulting in massive splenic B lymphocyte apoptosis through the mitochondrial pathway when compared to influenza infection alone^16^. However, these studies have been performed only in the context of superinfection, where alveolar macrophages are already depleted by virus infection. Coinfected animals showed the strongest Th1 skewed humoral response, which might be helping in the early viral clearance detected at 3DPI. In human clinical trials, while nasal pneumococcal colonization had no impact upon TIV-induced antibody responses in serum, pneumococcal colonization dampened vaccine-mediated mucosal antibody responses, measured as Influenza-binding IgA and IgG in nasal washes^67^. These results warrant further investigation of mucosal immune responses after challenge in our models.

Overall, our results highlight the importance of influenza vaccination in the prevention of severe disease following bacterial secondary infections. They also underline important differential outcomes of the antiviral host response during coinfections and superinfection and their divergent effects in antibody response to TIV vaccination. We show that both coinfection and superinfection cause neutrophilic inflammation in the lung that correlates with detrimental outcomes and is partially prevented by vaccination. However, SiglecF^hi^ neutrophils are increased in vaccinated animals and may play a protective role. More mechanistic studies are needed to understand the underlying immune mechanisms that drive these observations. Finally, we observed lung eosinophilia that correlated with protection from coinfection and superinfection in vaccinated animals, thereby confirming observations reported before by our research group for the influenza vaccination/challenge model.

## MATERIALS AND METHODS

### Ethics statement

All experiments were approved and carried out in compliance with the Institutional Biosafety Committee (IBC) and Institutional Animal Care and Use Committee (IACUC) regulations of Icahn School of Medicine at Mt. Sinai. Protocol number: IACUC-2017-0330

### Cells

MDCK cells (female origin Madin-Darby canine kidney cells, ATCC CRL-3216) were used to determine the NC99 titration. The cell lines were maintained in Dulbecco’s modified Eagle’s medium (DMEM, GIBCO) supplemented with 10% (v/v) fetal bovine serum (GIBCO) and 1x penicillin/streptomycin at 37°C under a humidified atmosphere containing 5% CO_2_.

### Bacteria

*Streptococcus pneumoniae* ATCC6301® (serotype 1) was grown and maintained at 37 °C, 5% CO2 on Columbia blood agar plates (Thermo Fisher). Overnight cultures were prepared by adding a single colony in 40 mL of brain heart infusion (BHI) broth (Sigma). Then, 3 mL of grown culture was transferred to a new tube with 12 mL of fresh BHI media and incubated between 4-6 h until it reached an Optical Density of 600nm (OD600) equal to 1. The number of bacteria in the final suspension was determined by plating 10-fold serial dilutions onto agar plates. Titration of bacteria showed that an OD600 approximately had 2.665×10^8^ CFU/ml. Then, 1 mL was centrifuged at 2000 rpms for 5 min. The supernatant was discarded, and the pellet was resuspended in 1 mL of PBS and diluted to obtain the desired number of bacteria for infection. Bacteria were identified as pneumococci by alpha-hemolysis on blood agar. Storage of bacteria was performed at mid-exponential phase (5-6 h) in BHI, with the addition of 10% glycerol, and storage at -80 °C. To enhance bacterial specificity to murine hosts, strains were passaged once within murine lungs, subsequently recovered, and cryopreserved according to the previously described protocol.

### Virus

The virus used for challenge was H1N1 A/New Caledonia/20/1999 (NC99) at a sublethal dose (0.4 LD_50_, 632.4 PFU/mL).

### Vaccine

For all vaccination experiments 1/5 of the human dose of the seasonal trivalent inactivated influenza virus vaccine was administered intramuscularly in the hind legs (TIV; Fluzone® Influenza Virus Vaccine) containing an influenza A H1N1 component (A/New Caledonia/20/1999/IVR-116), influenza A H3N2 component (A/New York/55/2004/X-157 [an A/California/7/2004-like strain]), and influenza B component (B/Jiangsu/10/2003 [a B/Shanghai/361/2002-like strain]).

### Mouse Infection Models

All experiments were performed with 6–8-week-old female C57BL/6 mice (Strain #000664, The Jackson Laboratory).

A standardized suspension of the ATCC6301® strain was prepared for inoculation by overnight growth in BHI from a single colony. The next day, subculture was performed, and the bacteria allowed to grow to mid-exponential phase (5-6 h). To standardize the initial dose of infection, optical density (OD600) was measured at different timepoints until the absorbance at 600nm reached values of 1. Then, bacteria were harvested by centrifugation and suspended in phosphate-buffered saline (PBS). This suspension was diluted 1/10 and titrated in Columbia blood agar. For the initial infection 1000 CFUs of ATCC6301® strain were used according to titrated values at OD600=1. For viral challenge 0.4 LD_50_ of NC99 was diluted in PBS to the appropriate infection volume.

Mice were intranasally challenged under mild ketamine/xylazine anesthesia. A final infection volume of 50 μL was used for all experiments. If coinfection was performed, bacteria and virus were combined in a single suspension administered in the same final volume. Detailed experimental layout is presented in Fig. 1a.

### Bacterial titatrion by Miles & Misra quantification

The upper lobe from the right lung of the mice was collected in 500 µL of 1X PBS. On the same day of the assay, lungs were disrupted using a cell strainer (70 µm) and the back of a sterile syringe. Homogenates were collected in 1.5 mL Eppendorf tubes. Serial 1/10 dilutions of each lung were performed in 96-well plates. Dilutions were plated by adding a single drop of 20 µL on Columbia blood agar plates according to Miles & Misra titration method^68^. Technical triplicates of each lung were performed. Agar plates were incubated overnight in an incubator at 37°C with 5% CO_2_. The next day, single colonies were counted to determine lung bacterial titers.

### Virus titration by plaque assays

NC99 titration was performed as previously described^45^. Briefly, the middle lobe from the right lung was collected in 500 µL of PBS in prefilled homogenizer bead tubes containing 3.0 mm high impact zirconium beads (Benchmark Scientific), snap-frozen on dry ice after harvest and stored at -80°C. On the day of the assay, samples were thawed, homogenized, and then centrifuged at 5,000 x g for 5 min at 4°C. Supernatants were 10-fold serially diluted in PBS from 1/10 to 1/10^6^. 12-well tissue culture plates were seeded the day prior with 3.5×10^5^ MDCK cells. The day of the assay, plates were washed once with 1 mL of PBS per well. After washing, 250 µL of each lung supernatant dilution was added to the plate and then incubated for 1 hour at room temperature (RT). After incubation, cells were washed once with 1 mL/well of PBS then 1 mL/well of overlay was added to each well. The overlay solidified at RT for 15 min and then plates were incubated at 37°C, 5% CO_2_ for 48 hr. Then, plates were fixed with 1 mL of 4% formalin per well O/N at 4°C. After washing with PBS-T (PBS+0.05% Tween 20), monolayers were immunostained with 200 μL/well polyclonal mouse serum diluted in blocking buffer (5% milk in PBS) for 1.5 hr on a plate rocker. Plates were washed 1 time with PBS-T then 200 μL/well of horseradish peroxidase (HRP)-conjugated goat anti-mouse IgG Fc secondary antibody diluted in blocking buffer for 1 hr at room temperature while rocking. Plates were washed, then 150 μL/well KPL TrueBlue was added and incubated at room temperature for 30 min on a plate rocker. Plates were washed and plaques were counted to determine lung viral titers.

### Flow cytometry

To analyze the lung cell populations at 6 and 10 DPI, the left lung lobe was collected and made into a single cell suspension using a Mouse Lung Dissociation Kit (Miltenyi Biotec), following manufacturer instructions. In brief, gentleMACS C tubes were filled with 2.4 mL of Buffer S and kept on ice while lungs were harvested. Then, each tube was spiked with 100 µL of Enzyme D, and 15 µL of Enzyme A. Then, C Tubes were attached it upside down onto the sleeve of the gentleMACS dissociator, where they were incubated at 37°C during 32 min, following the 37C_m_LDK_1 program of the gentleMACS dissociator.

After generating a single cell suspension, cells were centrifuged for 5mins at 500 x g and the supernatant was decanted. Then, cells were resuspended into 2 mL of RBC lysis solution (5 PRIME) and incubated at RT for 10 min. Then 10 mL of PBS was added to stop lysis, and cells were centrifuged for 5 mins at 500 x g and the supernatant was decanted. Then cells were resuspended into 50 μL of Fc Block and incubated for 5 mins at room temperature. Meanwhile a staining cocktail solution was prepared diluting 1 μL of BD Pharmingen™ FITC Hamster Anti Mouse CD3e (Clone 145-2C11), BD Pharmingen™ PE Rat Anti-CD11b (Clone M1/70), BD Horizon™ PE-CF594 Rat Anti-Mouse Siglec-F (Clone E50-2440), BD Pharmingen™ PE-Cy™7 Hamster Anti-Mouse CD11c (Clone HL3), Ly-6C Monoclonal Antibody, APC, eBioscience™, Invitrogen™ (Clone HK1.4),

Alexa Fluor® 700 anti-mouse Ly-6G Antibody (Clone 1A8) and 0.5 μL of Fixable Viability-e780 and MHC Class II (I-A/I-E) Monoclonal Antibody, eFluor™ 450, eBioscience™, (Clone M5/114.15.2); into a final volume of 50 μL in FACS buffer. The staining cocktail was added on top of the samples, and they were incubated for 20 mins at room temperature protected from the light. After the incubation period, cells were centrifuged for 5 mins at 500 x g and washed twice with 200 μL of FACS buffer. Samples were acquired using a Beckman Coulter Gallios flow cytometer with Kaluza software. Data analysis was performed using FlowJo v10.10 (Treestar) and compensated using the built-in AutoSpill algorithm. Data were visualized using GraphPad Prism v10.1.1.

### Lung cytokine and chemokine profiling

The middle lobe from the right lung of the mice was collected, homogenized, and clarified as described above. The Cytokine & Chemokine 26-Plex Mouse ProcartaPlex™ Panel 1 (ThermoFisher) was used to measure cytokine and chemokine concentrations in the lungs, following manufacturer’s instructions. In brief, 50 µL of lung homogenate was combined with beads in a flat-bottom black 96-well plate and incubated for 30 minutes at RT protected from light. All RT incubations were performed in a shaker at 300 rpm. After 30 mins, the plate was moved to 4°C for overnight incubation. The next day, the plate was equilibrated to room temperature for 30 ms, then washed 3 times with 150 µL/well of Wash Buffer. Following washing, 25 µL/well of Detection Antibody mixture was added and incubated at room temperature for 30 minutes. The plate was washed 3 times, then, 50 µL/well of Streptavidin-PE solution was added and incubated for 30 mins at RT. After washing the plate 3 times, 120 µL/well of Reading Buffer was added and the plate was incubated for 5 minutes at room temperature. Data were acquired on a Lumine× 100/200 analyzer (Millipore) with xPONENT software [version (v) 4.3]. Data visualization and analysis were conducted using GraphPad Prism (v10.1.1) and RStudio (v2022.12.0+353).

### Enzyme-linked Immunosorbent Assay (ELISA)

Mice sera were tested for vaccine-specific IgG, IgG1 and IgG2c titers. ELISA plates (Nunc MAXISORP, ThermoFisher) were coated with TIV vaccine diluted 1:100 into ELISA coating buffer (0.2M bicarbonate buffer, pH=9.6) and incubated O/N at 4°C. Plates were washed three times with washing buffer [1X PBS + 0.1% Tween20 (Sigma Aldrich)] and incubated in blocking buffer for 1 hour at room temperature. The serum samples were 3-fold serially diluted starting with 1:100 dilution in blocking buffer and allowed to bind to the TIV-coated ELISA plates for 1hr 30min. Then, the plates were washed three times with washing buffer and incubated with 1/5000 diluted secondary total IgG (Sigma Aldrich) or 1/4000 IgG1 or IgG2c antibodies (Southern Biotech), conjugated to horse radish peroxidase (HRP) for 1hr at room temperature. Finally, the plates were washed three times in washing buffer and incubated with 50µl of tetramethylbenzidine substrate (TMB, BD) until blue color appeared. The colorimetric reaction was stopped by adding 50µl of ELISA stop solution (ThermoFisher) and absorbance was measured at 450nm and 650nm wavelengths.

### Statistics

Pairwise comparisons were made by Mann-Whitney U test. Differences among three or more groups were performed by One-Way ANOVA with Tukey’s multiple comparisons test and survival was assessed by Log Rank (Mantel-Cox) test. Statistical test is specified in each case in the figure legend. Significance values are represented as * p≤0.05, ** p≤0.01, *** p≤0.001, **** p≤0.0001. GraphPad Prism (v10.1.1) and RStudio (v2022.12.0+353) were used for analysis and visualizations.

## Acknowledgements

We thank the Mount Sinai Center for Comparative Medicine and Surgery for mouse colony management.

## SUPPLEMENTARY FIGURES

**Supplementary Figure 1.**
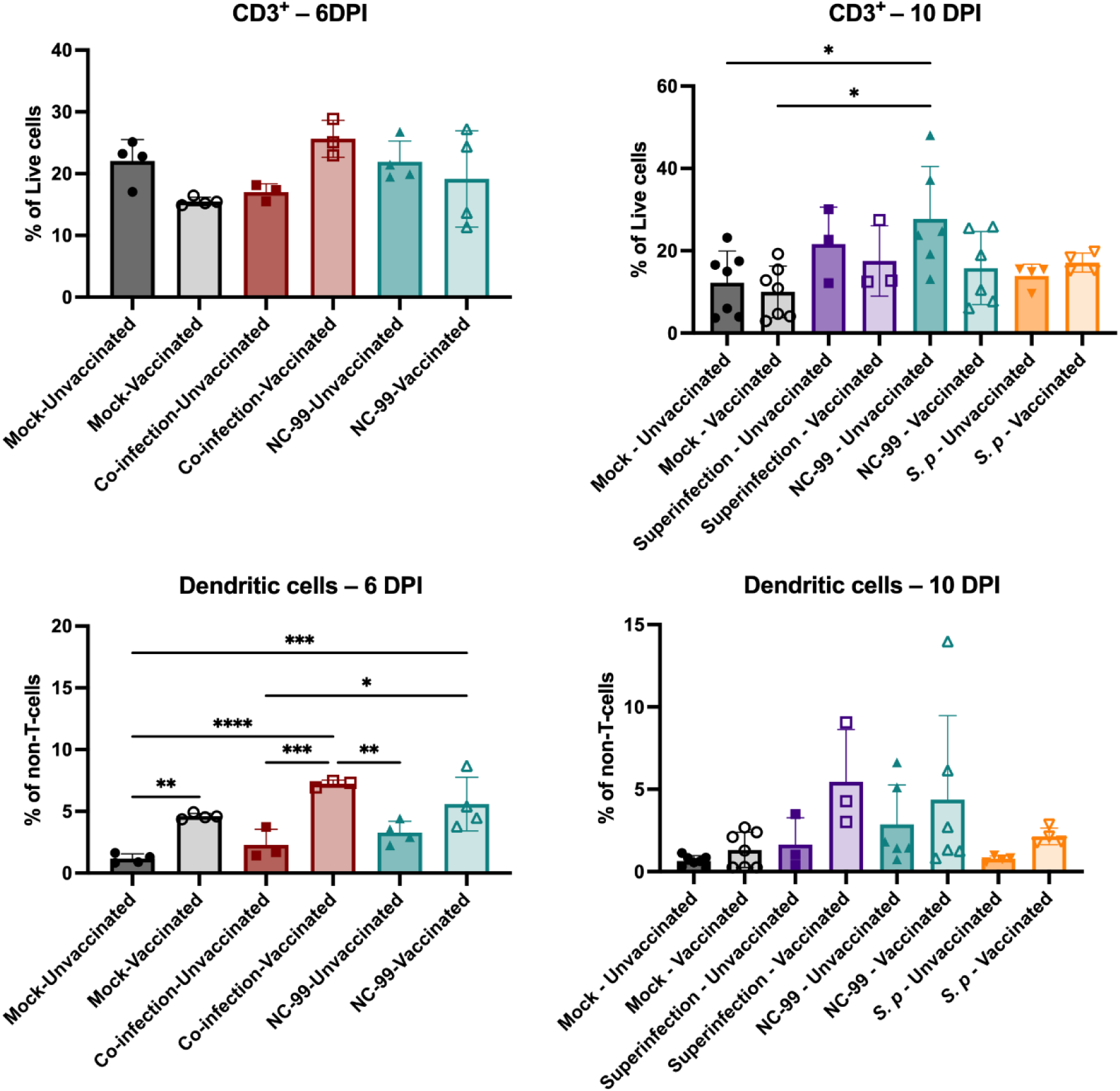
Relative quantification of lung CD3^+^ and dendritic cells at 6 and 10 DPI. Comparisons were performed by One-Way ANOVA with Tukey’s multiple comparisons test. Significance values are represented as * p≤0.05, ** p≤0.01, *** p≤0.001, **** p≤0.0001.

**Supplementary Figure 2.**
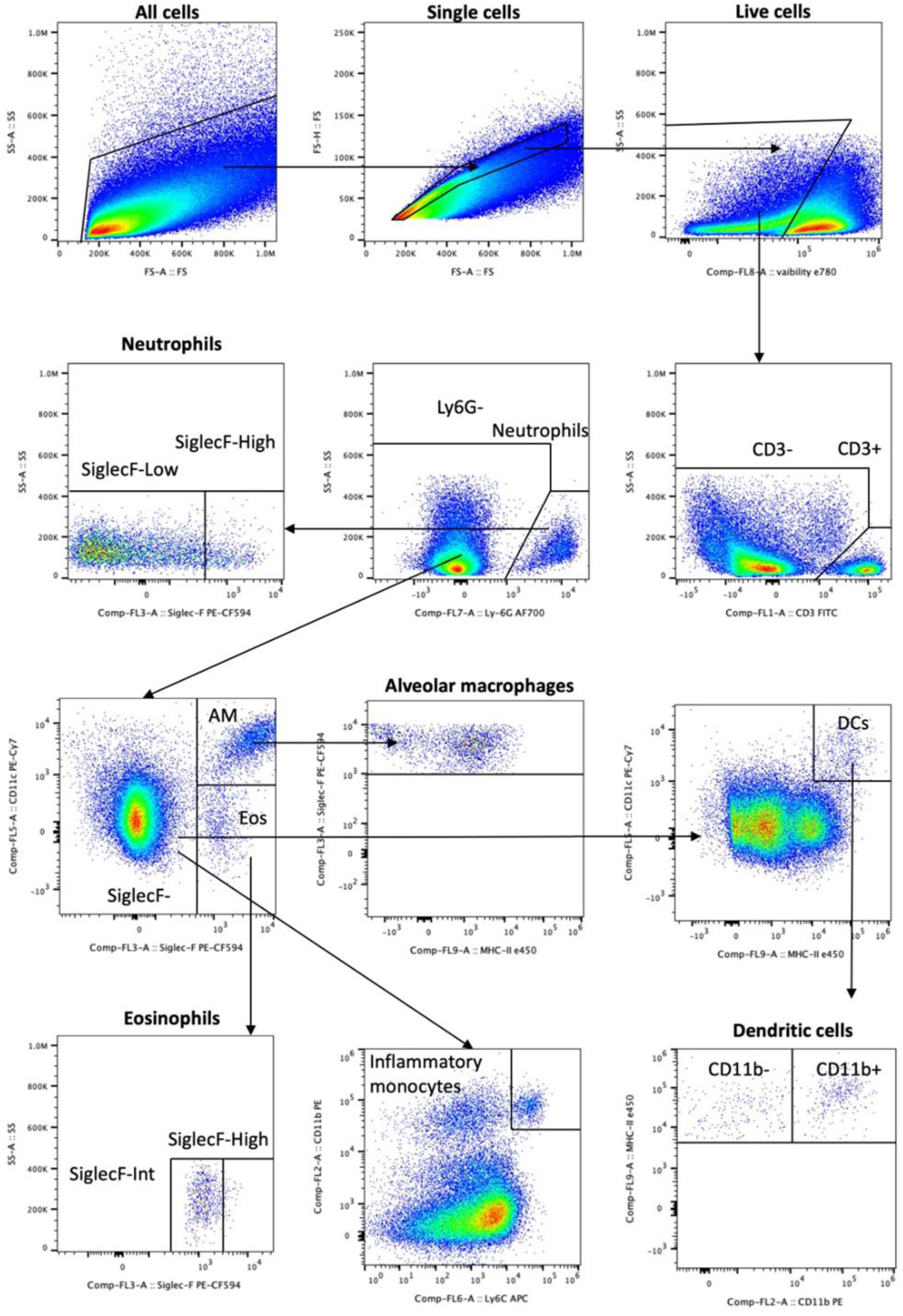
Gating strategy for classification of lung cell populations. Gating strategy, exemplified in an influenza NC99 infected lung at 10 DPI.

**Supplementary Figure 3.**
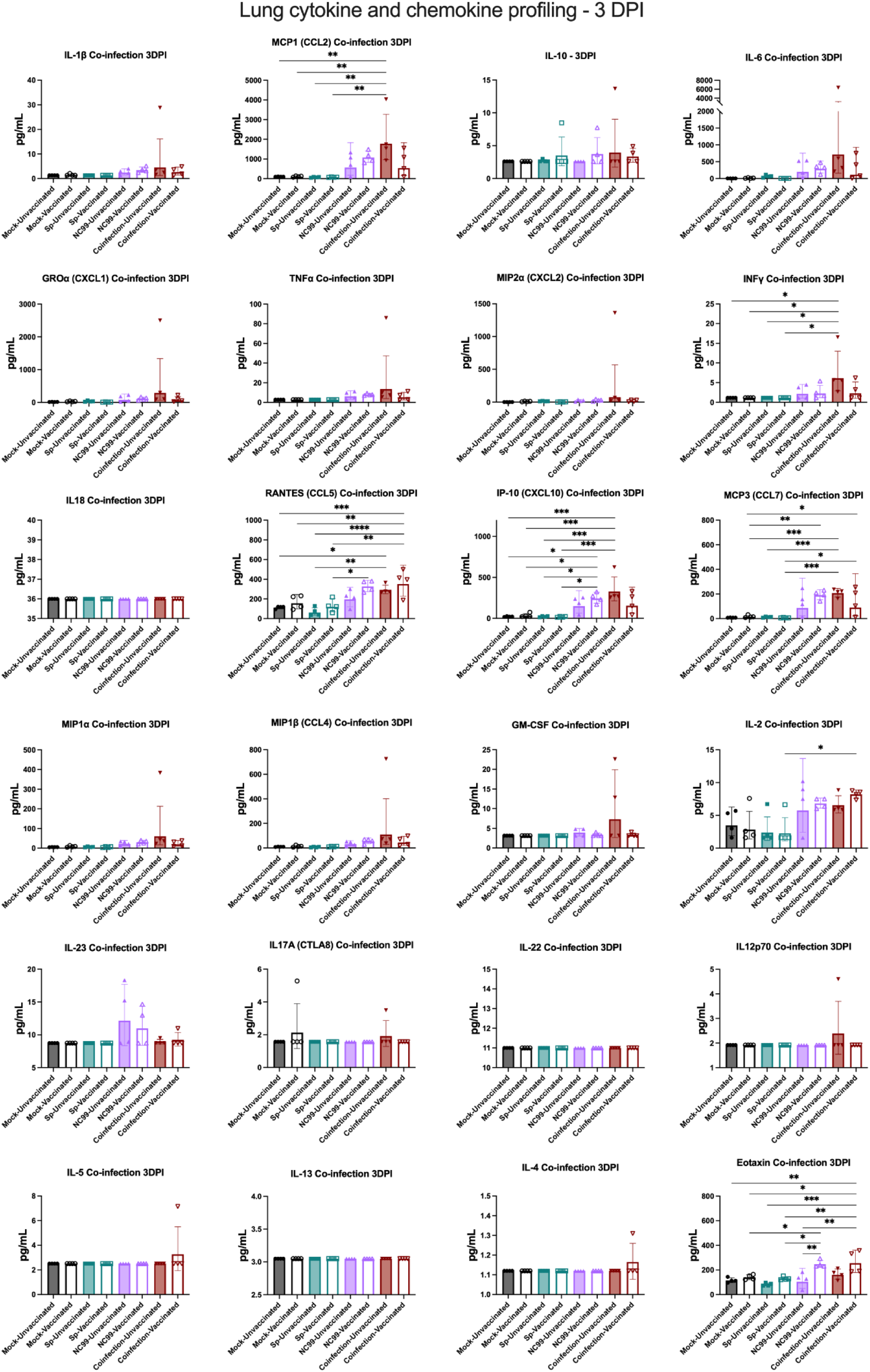
lung homogenate supernatant cytokine/chemokine data at 3 DPI. Bar graphs (Mean±SD) of absolute concentration (pg/ml) of cytokines and chemokines in lungs.

**Supplementary Figure 4.**
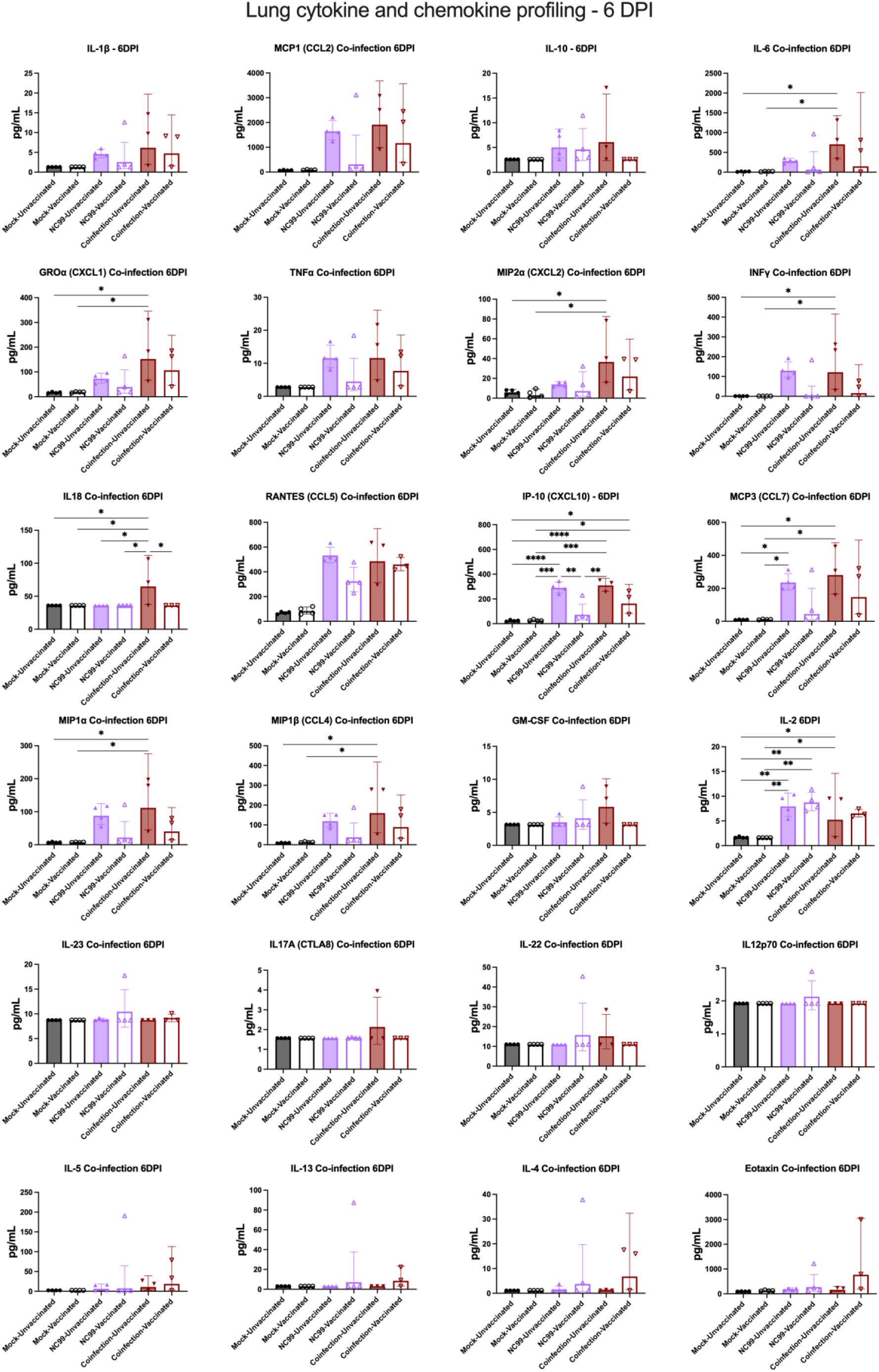
lung homogenate supernatant cytokine/chemokine data at 6 DPI. Bar graphs (Mean±SD) of absolute concentration (pg/ml) of cytokines and chemokines in lungs.

**Supplementary Figure 5.**
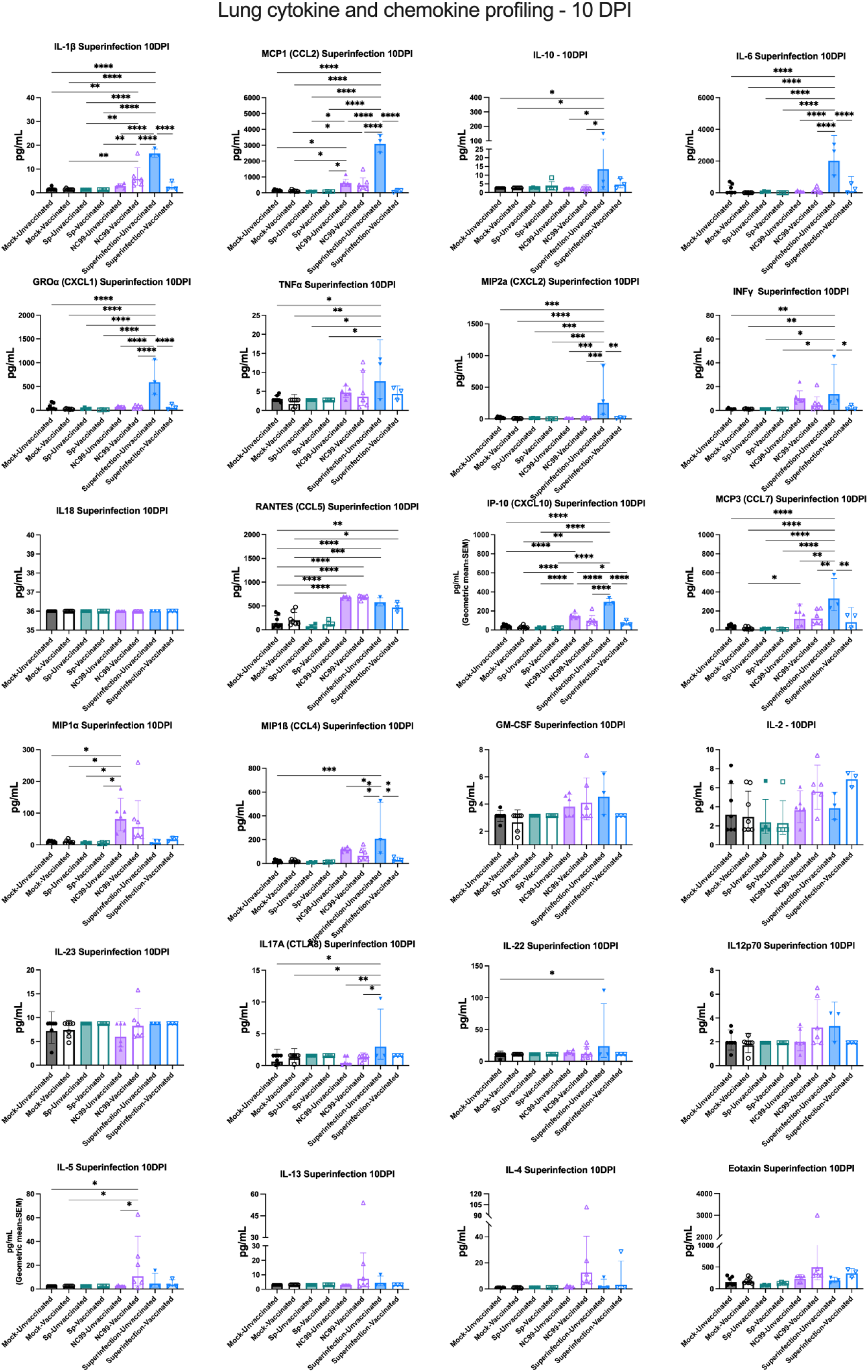
lung homogenate supernatant cytokine/chemokine data at 10 DPI. Bar graphs (Mean±SD) of absolute concentration (pg/ml) of cytokines and chemokines in lungs.

## REFERENCES

1. Morris, D. E., Cleary, D. W. & Clarke, S. C. Secondary Bacterial Infections Associated with Influenza Pandemics. Front. Microbiol. 8, (2017).

2. Morens, D. M., Taubenberger, J. K. & Fauci, A. S. Predominant Role of Bacterial Pneumonia as a Cause of Death in Pandemic Influenza: Implications for Pandemic Influenza Preparedness. J. Infect. Dis. 198, 962–970 (2008).

3. Hers, J. F. P., Masurel, N. & Mulder, J. BACTERIOLOGY AND HISTOPATHOLOGY OF THE RESPIRATORY TRACT AND LUNGS IN FATAL ASIAN INFLUENZA. The Lancet 272, 1141–1143 (1958).

4. Louria, D. B., Blumenfeld, H. L., Ellis, J. T., Kilbourne, E. D. & Rogers, D. E. STUDIES ON INFLUENZA IN THE PANDEMIC OF 1957-1958. II. PULMONARY COMPLICATIONS OF INFLUENZA*†. J. Clin. Invest. 38, 213–265 (1959).

5. Iuliano, A. D. et al. Estimates of global seasonal influenza-associated respiratory mortality: a modelling study. The Lancet 391, 1285–1300 (2018).

6. McCullers, J. A. The co-pathogenesis of influenza viruses with bacteria in the lung. Nat. Rev. Microbiol. 12, 252–262 (2014).

7. Arranz-Herrero, J. et al. Determinants of poor clinical outcome in patients with influenza pneumonia: A systematic review and meta-analysis. Int. J. Infect. Dis. 131, 173–179 (2023).

8. Weis, S. & Te Velthuis, A. J. W. Influenza Virus RNA Synthesis and the Innate Immune Response. Viruses 13, 780 (2021).

9. Mifsud, E. J., Kuba, M. & Barr, I. G. Innate Immune Responses to Influenza Virus Infections in the Upper Respiratory Tract. Viruses 13, 2090 (2021).

10. Klomp, M., Ghosh, S., Mohammed, S. & Nadeem Khan, M. From virus to inflammation, how influenza promotes lung damage. J. Leukoc. Biol. 110, 115–122 (2021).

11. Iverson, A. R. et al. Influenza Virus Primes Mice for Pneumonia From Staphylococcus aureus. J. Infect. Dis. 203, 880–888 (2011).

12. Hanada, S., Pirzadeh, M., Carver, K. Y. & Deng, J. C. Respiratory Viral Infection-Induced Microbiome Alterations and Secondary Bacterial Pneumonia. Front. Immunol. 9, (2018).

13. Smith, A. M. et al. Kinetics of Coinfection with Influenza A Virus and Streptococcus pneumoniae. PLoS Pathog. 9, e1003238 (2013).

14. Smith, A. P. et al. Dynamic Pneumococcal Genetic Adaptations Support Bacterial Growth and Inflammation during Coinfection with Influenza. Infect. Immun. 89, (2021).

15. Karwelat, D. et al. Influenza virus-mediated suppression of bronchial Chitinase-3-like 1 secretion promotes secondary pneumococcal infection. FASEB J. 34, 16432–16448 (2020).

16. Xiang, L. Influenza a virus and *Streptococcus pneumonia* coinfection potentially promotes bacterial colonization and enhances B lymphocyte depression and reduction. J. Biol. Regul. Homeost. Agents 33, 1437–1449 (2019).

17. Rowe, H. M. et al. Direct interactions with influenza promote bacterial adherence during respiratory infections. Nat. Microbiol. 4, 1328–1336 (2019).

18. Brundage, J. F. Interactions between influenza and bacterial respiratory pathogens: implications for pandemic preparedness. Lancet Infect. Dis. 6, 303–312 (2006).

19. Metzger, D. W. & Sun, K. Immune Dysfunction and Bacterial Coinfections following Influenza. J. Immunol. 191, 2047–2052 (2013).

20. McCullers, J. A. & Bartmess, K. C. Role of Neuraminidase in Lethal Synergism between Influenza Virus and*Streptococcus pneumoniae*. J. Infect. Dis. 187, 1000–1009 (2003).

21. Van Der Sluijs, K. F. et al. Involvement of the platelet-activating factor receptor in host defense against*Streptococcus pneumoniae*during postinfluenza pneumonia. Am. J. Physiol.-Lung Cell. Mol. Physiol. 290, L194–L199 (2006).

22. Nugent, K. M. & Pesanti, E. L. Tracheal function during influenza infections. Infect. Immun. 42, 1102–1108 (1983).

23. Verma, A. K., Bansal, S., Bauer, C., Muralidharan, A. & Sun, K. Influenza Infection Induces Alveolar Macrophage Dysfunction and Thereby Enables Noninvasive *Streptococcus pneumoniae* to Cause Deadly Pneumonia. J. Immunol. 205, 1601–1607 (2020).

24. McNamee, L. A. & Harmsen, A. G. Both Influenza-Induced Neutrophil Dysfunction and Neutrophil-Independent Mechanisms Contribute to Increased Susceptibility to a Secondary*Streptococcus pneumoniae*Infection. Infect. Immun. 74, 6707–6721 (2006).

25. van der Sluijs, K. F. et al. Influenza-Induced Expression of Indoleamine 2,3-Dioxygenase Enhances Interleukin-10 Production and Bacterial Outgrowth during Secondary Pneumococcal Pneumonia. J. Infect. Dis. 193, 214–222 (2006).

26. Van Der Sluijs, K. F. et al. IL-10 Is an Important Mediator of the Enhanced Susceptibility to Pneumococcal Pneumonia after Influenza Infection. J. Immunol. 172, 7603–7609 (2004).

27. Jakab, G. J. & Green, G. M. Defect in intracellular killing of Staphylococcus aureus within alveolar macrophages in Sendai virus-infected murine lungs. J. Clin. Invest. 57, 1533–1539 (1976).

28. Jakab, G. J., Warr, G. A. & Sannes, P. L. Alveolar Macrophage Ingestion and Phagosome-Lysosome Fusion Defect Associated with Virus Pneumonia. Infect. Immun. 27, 960–968 (1980).

29. Didierlaurent, A. et al. Sustained desensitization to bacterial Toll-like receptor ligands after resolutionof respiratory influenza infection. J. Exp. Med. 205, 323–329 (2008).

30. Zavitz, C. C. J. et al. Dysregulated Macrophage-Inflammatory Protein-2 Expression Drives Illness in Bacterial Superinfection of Influenza. J. Immunol. 184, 2001–2013 (2010).

31. Godfroid, F., Hermand, P., Verlant, V., Denoël, P. & Poolman, J. T. Preclinical Evaluation of the Pht Proteins as Potential Cross-Protective Pneumococcal Vaccine Antigens. Infect. Immun. 79, 238–245 (2011).

32. Pan, W. et al. Visualizing influenza virus infection in living mice. Nat. Commun. 4, (2013).

33. Califano, D., Furuya, Y. & Metzger, D. W. Effects of Influenza on Alveolar Macrophage Viability Are Dependent on Mouse Genetic Strain. J. Immunol. 201, 134–144 (2018).

34. Gouma, S., Anderson, E. M. & Hensley, S. E. Challenges of Making Effective Influenza Vaccines. Annu. Rev. Virol. 7, 495–512 (2020).

35. Wei, C.-J. et al. Next-generation influenza vaccines: opportunities and challenges. Nat. Rev. Drug Discov. 19, 239–252 (2020).

36. Grijalva, C. G. et al. Association Between Hospitalization With Community-Acquired Laboratory-Confirmed Influenza Pneumonia and Prior Receipt of Influenza Vaccination. JAMA 314, 1488 (2015).

37. Tessmer, A. et al. Influenza vaccination is associated with reduced severity of community-acquired pneumonia. Eur. Respir. J. 38, 147–153 (2011).

38. Choi, A., Christopoulou, I., Saelens, X., García-Sastre, A. & Schotsaert, M. TIV Vaccination Modulates Host Responses to Influenza Virus Infection that Correlate with Protection against Bacterial Superinfection. Vaccines 7, 113 (2019).

39. Zurli, V. et al. Positive Contribution of Adjuvanted Influenza Vaccines to the Resolution of Bacterial Superinfections. J. Infect. Dis. 213, 1876–1885 (2016).

40. Okamoto, S. et al. Vaccination with formalin-inactivated influenza vaccine protects mice against lethal influenza Streptococcus pyogenes superinfection. Vaccine 22, 2887–2893 (2004).

41. Ritchie, N. D., Mitchell, T. J. & Evans, T. J. What is Different About Serotype 1 Pneumococci? Future Microbiol. 7, 33–46 (2012).

42. Wang, H. & Bozinovski, S. Emergence of an immunosuppressive neutrophil phenotype during pneumococcal lung co-infection with influenza A virus. in Airway cell biology and immunopathology PA2377 (European Respiratory Society, 2019). doi:10.1183/13993003.congress-2019.PA2377.

43. Chang, L. A. & Schotsaert, M. Ally, adversary, or arbitrator? The context-dependent role of eosinophils in vaccination for respiratory viruses and subsequent breakthrough infections. J. Leukoc. Biol. 116, 224–243 (2024).

44. Ebenig, A. et al. Vaccine-associated enhanced respiratory pathology in COVID-19 hamsters after TH2-biased immunization. Cell Rep. 40, 111214 (2022).

45. Chang, L. A. et al. Influenza breakthrough infection in vaccinated mice is characterized by non-pathological lung eosinophilia. Front. Immunol. 14, 1217181 (2023).

46. Diego, J. G.-B. et al. Breakthrough infections by SARS-CoV-2 variants boost cross-reactive hybrid immune responses in mRNA-vaccinated Golden Syrian hamsters. PLOS Pathog. 20, e1011805 (2024).

47. Mesnil, C. et al. Lung-resident eosinophils represent a distinct regulatory eosinophil subset. J. Clin. Invest. 126, 3279–3295 (2016).

48. Azzoni, R. et al. Distinct eosinophil subsets are modulated by agonists of the commensal-metabolite and vitamin B3 receptor GPR109A during allergic-type inflammation. Preprint at 10.1101/2022.08.04.502285 (2022).

49. Choi, A. et al. Non-sterilizing, Infection-Permissive Vaccination With Inactivated Influenza Virus Vaccine Reshapes Subsequent Virus Infection-Induced Protective Heterosubtypic Immunity From Cellular to Humoral Cross-Reactive Immune Responses. Front. Immunol. 11, 1166 (2020).

50. Ghoneim, H. E., Thomas, P. G. & McCullers, J. A. Depletion of Alveolar Macrophages during Influenza Infection Facilitates Bacterial Superinfections. J. Immunol. 191, 1250–1259 (2013).

51. Wu, N.-H., Meng, F., Seitz, M., Valentin-Weigand, P. & Herrler, G. Sialic acid-dependent interactions between influenza viruses and Streptococcus suis affect the infection of porcine tracheal cells. J. Gen. Virol. 96, 2557–2568 (2015).

52. Wang, Y. et al. Capsular Sialic Acid of Streptococcus suis Serotype 2 Binds to Swine Influenza Virus and Enhances Bacterial Interactions with Virus-Infected Tracheal Epithelial Cells. Infect. Immun. 81, 4498–4508 (2013).

53. King, S. J., Hippe, K. R. & Weiser, J. N. Deglycosylation of human glycoconjugates by the sequential activities of exoglycosidases expressed by *Streptococcus pneumoniae*. Mol. Microbiol. 59, 961–974 (2006).

54. McCullers, J. A. & Rehg, J. E. Lethal Synergism between Influenza Virus and *Streptococcus pneumoniae:* Characterization of a Mouse Model and the Role of Platelet-Activating Factor Receptor. J. Infect. Dis. 186, 341–350 (2002).

55. Baker, P. J. et al. Co-infection of mice with SARS-CoV-2 and Mycobacterium tuberculosis limits early viral replication but does not affect mycobacterial loads. Front. Immunol. 14, 1240419 (2023).

56. Aegerter, H. et al. Influenza-induced monocyte-derived alveolar macrophages confer prolonged antibacterial protection. Nat. Immunol. 21, 145–157 (2020).

57. Damjanovic, D., Lai, R., Jeyanathan, M., Hogaboam, C. M. & Xing, Z. Marked Improvement of Severe Lung Immunopathology by Influenza-Associated Pneumococcal Superinfection Requires the Control of Both Bacterial Replication and Host Immune Responses. Am. J. Pathol. 183, 868–880 (2013).

58. Tavares, L. P. et al. CXCR1/2 Antagonism Is Protective during Influenza and Post-Influenza Pneumococcal Infection. Front. Immunol. 8, 1799 (2017).

59. Shin, J. W. et al. A unique population of neutrophils generated by air pollutant– induced lung damage exacerbates airway inflammation. J. Allergy Clin. Immunol. 149, 1253–1269.e8 (2022).

60. Matsui, M., Nagakubo, D., Satooka, H. & Hirata, T. A novel Siglec-F+ neutrophil subset in the mouse nasal mucosa exhibits an activated phenotype and is increased in an allergic rhinitis model. Biochem. Biophys. Res. Commun. 526, 599–606 (2020).

61. Borkner, L., Curham, L. M., Wilk, M. M., Moran, B. & Mills, K. H. G. IL-17 mediates protective immunity against nasal infection with Bordetella pertussis by mobilizing neutrophils, especially Siglec-F+ neutrophils. Mucosal Immunol. 14, 1183–1202 (2021).

62. Prince, G. A. et al. Enhancement of respiratory syncytial virus pulmonary pathology in cotton rats by prior intramuscular inoculation of formalin-inactiva ted virus. J. Virol. 57, 721–728 (1986).

63. Waris, M. E., Tsou, C., Erdman, D. D., Zaki, S. R. & Anderson, L. J. Respiratory synctial virus infection in BALB/c mice previously immunized with formalin-inactivated virus induces enhanced pulmonary inflammatory response with a predominant Th2-like cytokine pattern. J. Virol. 70, 2852–2860 (1996).

64. Drake, M. G. et al. Human and Mouse Eosinophils Have Antiviral Activity against Parainfluenza Virus. Am. J. Respir. Cell Mol. Biol. 55, 387–394 (2016).

65. Percopo, C. M. et al. Activated mouse eosinophils protect against lethal respiratory virus infection. Blood 123, 743–752 (2014).

66. Wu, Y. et al. Lethal Coinfection of Influenza Virus and Streptococcus pneumoniae Lowers Antibody Response to Influenza Virus in Lung and Reduces Numbers of Germinal Center B Cells, T Follicular Helper Cells, and Plasma Cells in Mediastinal Lymph Node. J. Virol. 89, 2013–2023 (2015).

67. Carniel, B. F. et al. Pneumococcal colonization impairs mucosal immune responses to Live Attenuated Influenza Vaccine in adults. JCI Insight (2021) doi:10.1172/jci.insight.141088.

68. Miles, A. A., Misra, S. S. & Irwin, J. O. The estimation of the bactericidal power of the blood. Epidemiol. Infect. 38, 732–749 (1938).

